# New paradigms in actomyosin energy transduction: critical evaluation of non-traditional models for orthophosphate release

**DOI:** 10.1101/2023.04.12.535196

**Authors:** Alf Månsson, Marko Usaj, Luisa Moretto, Oleg Matusovsky, Lok Priya Velayuthan, Ran Friedman, Dilson E Rassier

## Abstract

Release of the ATP hydrolysis product inorganic phosphate (Pi) from the active site of myosin is central in chemo-mechanical energy transduction and closely associated with the main force-generating structural change, the power-stroke. Despite intense investigations, the relative timing between Pi-release and the power-stroke remains poorly understood. This hampers in depth understanding of the production of force and motion by myosin in health and disease and also our understanding of myosin-active drugs. From the 1990s and up to today, models with the Pi-release either distinctly before or after the power-stroke, in unbranched kinetic schemes, have dominated the literature. However, in recent years, alternative models have emerged to explain apparently contradictory findings. Here, we first compare and critically analyze, three influential alternative models, either characterized by a branched kinetic scheme or by partial uncoupling of Pi-release and the power-stroke. Finally, we suggest critical tests of the models aiming for a unified picture.

## Introduction

Adenosine triphosphate (ATP)-powered generation of force and motion by myosin molecules on actin filaments is central in eukaryotes [1–3]. The contractile machinery, consisting of actin, myosin and accessory proteins, is particularly well-organized in striated muscle (skeletal muscle and heart) with the proteins assembled in µm wide myofibrils that fill the muscle cells (Fig. 1A). The myofibrils consist of series-connected half-sarcomeres which, in turn, are composed of myosin-containing thick filaments and overlapping actin-containing thin filaments in a regular lattice (Fig. 1B). The main focus in this paper is myosin II (for its key structural elements, see Fig. 1C). The molecule has a tail domain that is the basis for filament formation and two motor domains (subfragments 1; S1). Each motor domain consists of a catalytic domain with an actin-binding region and an active site for ATP turnover (Fig. 1D). It also has an α-helical extension that is stabilized by one essential and one regulatory light chain allowing it to act as a lever arm (Fig. 1D). This lever arm swings during the force- and motion-generating structural change [2], the “power stroke”. The power stroke is closely associated with release of the ATP hydrolysis product orthophosphate (or inorganic phosphate; Pi) [4–25] but the exact temporal relationship between the Pi-release and the power stroke has been a long-standing enigma [7,10,12,15,16,21,24,26–29]. It is also unclear exactly what happens when externally added Pi-rebinds to myosin [18,27,30–34].

**Fig. 1.**
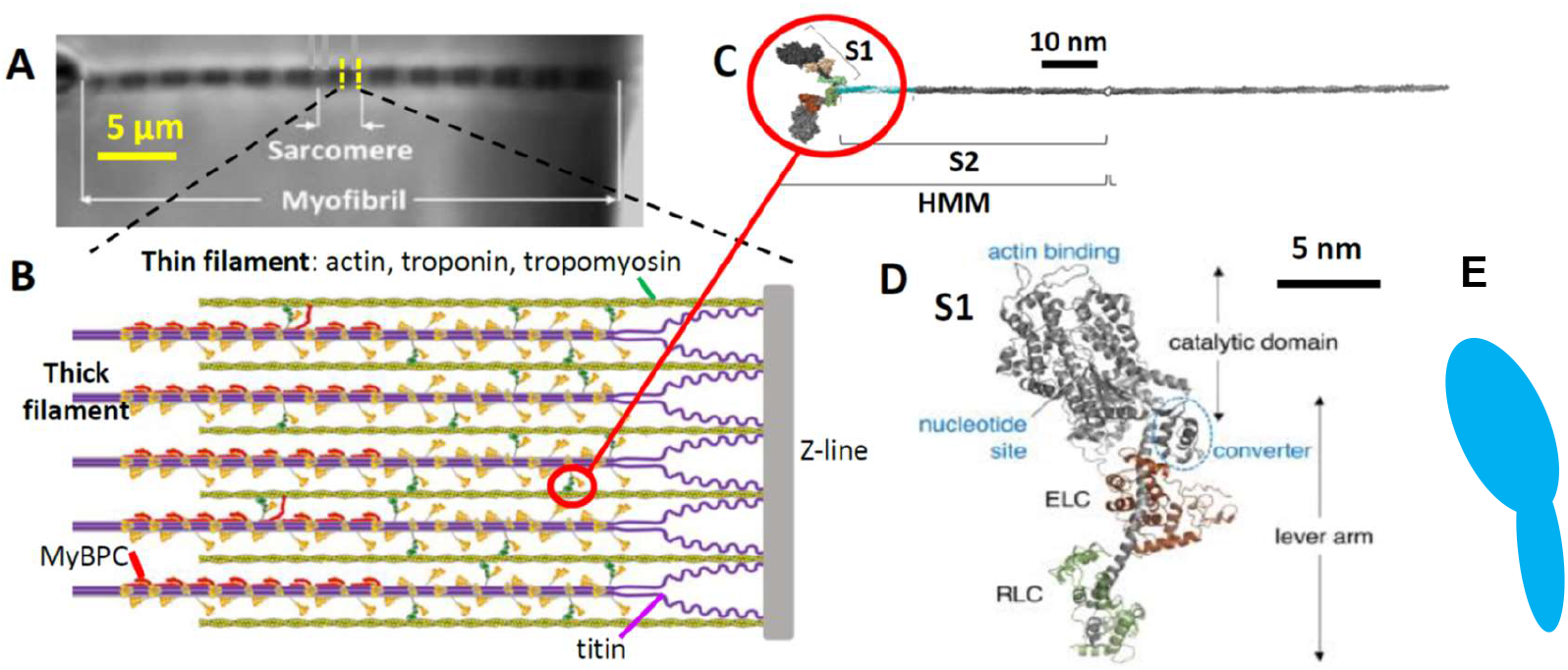
Striated muscle as example of actin-myosin contractile system. **A.** An isolated myofibril, mounted for force-measurements [35]. Myofibrils fill the muscle cells and are made up of repetitive structures, sarcomeres. **B.** Schematic of half-sarcomere with overlapping, thick, myosin-containing filaments and thin, actin-containing filaments (with regulatory proteins troponin and tropomyosin) attached to Z-line. Accessory proteins, myosin binding protein C (MyBPC) and titin also shown. **C.** Myosin molecule with two motor domains (heads or sub-fragment 1; S1). The proteolytic fragment heavy meromyosin (HMM) is also indicated. Note, that the orientation of the myosin molecule is flipped horizontally compared to its orientation in the half-sarcomere in B. **D.** Myosin motor domain with key functional elements indicated as well as essential (ELC) and regulatory (RLC) light chains. **E.** Schematic drawing of myosin S1 as used in figures below. Panels B-D from [36] reproduced under license CC BY 4.0.

Early models [37,38] assumed that the power-stroke was simultaneous with Pi-release. In contrast, more recent models have separated the power stroke and the Pi-release, placing Pi-release either distinctly before (Fig. 2A) or distinctly after (Fig. 2B) the power stroke [7,8,10,16,20]. The idea of Pi-release after the power stroke is consistent with transient biochemical kinetics studies combined with time-resolved fluorescence resonance energy transfer (FRET) used for detection of the power-stroke [7]. The idea is also in accordance with results evaluating transient mechanical responses to perturbations of length or tension at varied [Pi] in muscle cells and isolated myosin motors [6,10]. On the other hand, data from mechanical experiments on isolated myosin and actin by another group [39], as well as X-ray crystallography and reverse genetics studies [2,16] argue for Pi-release before the power-stroke. Similar conclusions have been reached based on theoretical analyses [28], implying that Pi-release after the power-stroke would cause reduced muscle shortening velocity and only a minimal change in isometric force upon increased [Pi], contrary to experimental findings [5,13,23,30]. The mentioned models, with Pi-release occurring either before or after the power-stroke without any branched pathways or other additional assumptions, are denoted “conventional models” below. This notation contrasts these models with recent “alternative models” that are the major focus here.

Recently, we built on ideas arising from experiments using myosin VI [2,16] to develop one such alternative model (Fig. 2C) where the power-stroke is partly uncoupled from Pi-release. This model assumes that, whereas Pi leaves the myosin active site before the power stroke, it pauses at secondary and tertiary sites on its way to bulk solution. This explains the observed delay between the power-stroke and Pi-appearance in solution [7,8,40,41]. Interestingly, this model also reconciles key contractile effects of varied [Pi], previously viewed as contradictory, including the [Pi]-independence of both shortening velocity and the power-stroke rate and a monotonous decrease in isometric force with increased [Pi]. Additionally, in our recent study [34], we used high-speed atomic force microscopy to verify the model prediction that the power-stroke can occur with only ADP (i.e. without Pi) at the myosin active site.

Despite the explanatory power of the described model with multistep Pi-release [34] (“multistep Pi-release model” below), outstanding issues remained. First, whereas we presented evidence for the existence of both secondary and tertiary Pi-binding sites in striated muscle myosin II we could not unequivocally demonstrate that the kinetics is slow enough to account for the observed rate of Pi-release (5-100 ms time delay) [7,16,41–43]. Moreover, we did not compare the multistep Pi-release model to other alternative models, e.g. with regard to conceptual similarities and/or if they would be as successful in accounting for a range of experimental data in a self-consistent way. This includes one model with a branched kinetic pathway (denoted “branched model” below) where Pi-release is assumed to occur after the power-stroke but where the cross-bridges may detach from a post-power-stroke state upon Pi-re-binding [18,27,30]. A third alternative model assumes loose coupling (“loose-coupling model” below) between the power-stroke and Pi-release (as well as ADP release) from the active site, where either the power-stroke or the Pi-release may occur first [11,44]. The latter model has similarities to the multistep Pi-release model [34] but does not include secondary and tertiary Pi-binding sites.

Here, we will first review the distinguishing features of the three alternative models. We will point out strengths and weaknesses of the different models and consider critical predictions and how these might be tested.

### Details of current models for Pi-release and re-binding

Fig. 2 describes current models relating the processes of Pi-release (yellow highlight in Fig. 2) and Pi-rebinding to the power-stroke (green highlight in Fig. 2). Mechanokinetic models, like those in Fig. 2, incorporate key molecular properties (strain-dependent actomyosin interaction kinetics, myosin elasticity and coarse grain structure) into kinetic schemes to allow quantitative predictions of contractile function [37,45–50]. The strain dependence is expressed in terms of a variable x that reflects the distance, along the actin and myosin filaments, between a myosin head and the nearest binding site on actin. This variable is also explicitly related to the strain of the cross-bridge elastic element, with different relationships for different biochemical cross-bridge states (see below). The use of mechanokinetic models enables rigorous tests of how different molecular mechanisms of Pi-release account for experimental data from single molecules, small ensembles of isolated actin and myosin or huge actin-myosin ensembles of muscle [45–47,50–52].

**Fig. 2.**
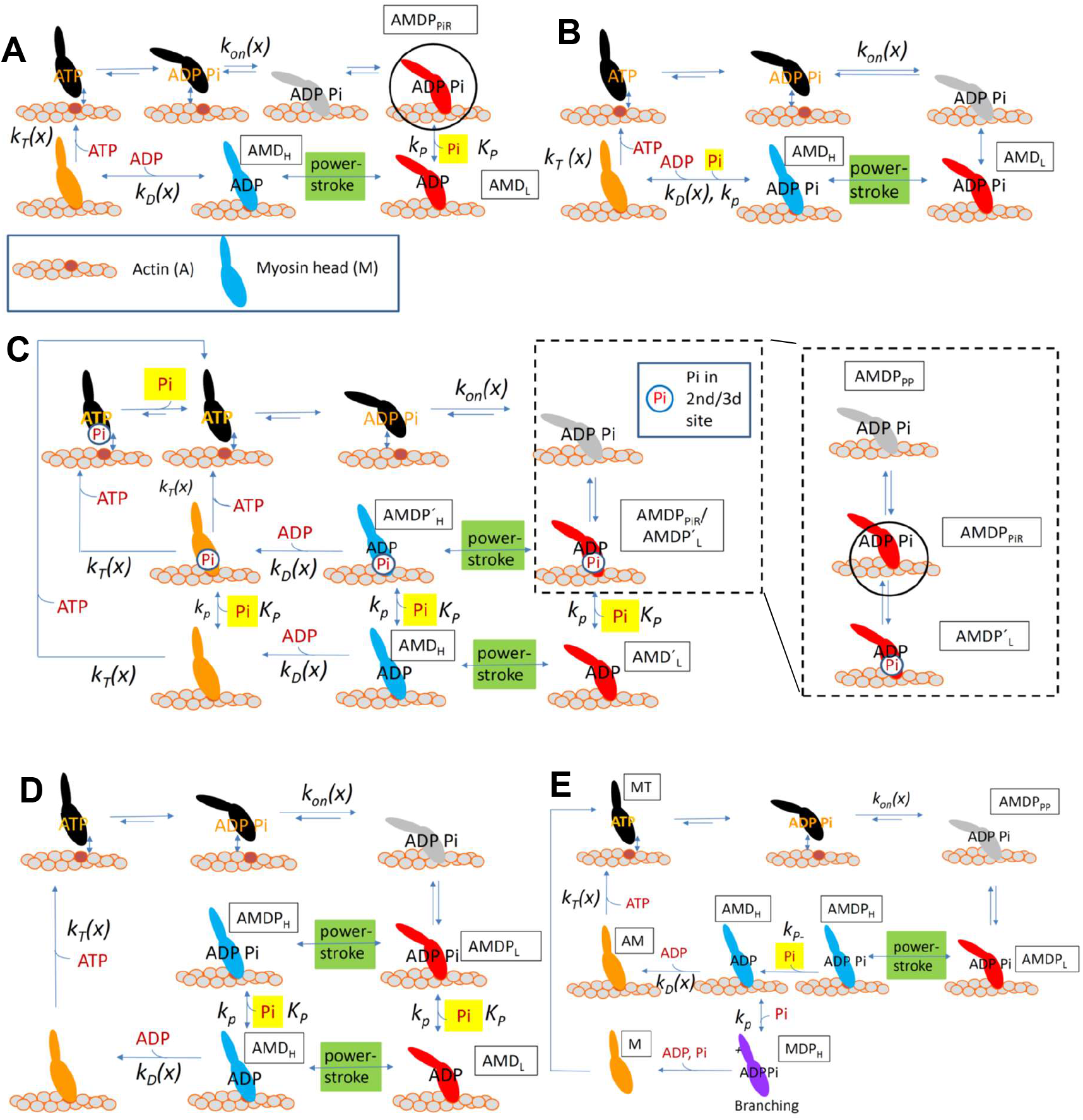
Mechanokinetic actomyosin cross-bridge models reflecting current ideas for relationship between force-generating power-stroke and the release of Pi from the myosin active site. **A.** Model with Pi-release from a Pi-release state (encircled) prior to force-generating power-stroke (from [20]). **B.** Model with Pi-release after power-stroke. **C.** Multistep Pi-release model from Moretto et al. [34] with Pi either in the active site (next to ADP in figure) or at a secondary/tertiary site (encircled Pi; full definition in text. Note, the rates along the second and third rows in the scheme are the same. Pi-release state (subscript PiR) is here lumped together with the AMDP_L_ start-of-powerstroke state. More details are given in inset within dashed box. **D.** Simplified version of loose-coupling model [11] focusing on the Pi-release and its temporal relationship to the power-stroke. It is assumed that Pi can be released from the active site either before or after the power-stroke, but, note, no secondary/tertiary Pi-binding site. **E.** Branched kinetic model [18,30,31] where Pi-rebinding leads to detachment into a post-power-stroke structural state (lower row). Key states in panels A-E, labelled by text boxes, with A: actin, M: myosin; T: ATP, D: ADP and P: inorganic phosphate. Rate constants and equilibrium constants are given by lower case and upper case letters, respectively. PiR, phosphate release state; H, high force; L, low force. The argument, (x), indicates that the rate constant (rate function) varies with the strain of the cross-bridge elastic element. Color coding: Detached states (black) and structurally different actin-attached states (different colors).

The models in Fig. 2 A and B, represent conventional models with Pi-release from the active site either distinctly before (e.g. [2,16,20,28,52,53] or after (e.g. [7,9,10,12,13,53]) the power-stroke, without any branched pathways. The models in Fig. 2 C-E, represent the alternative models, including the multistep Pi-release model of Moretto et al. [34] (Fig. 2 C), the loose coupling model of Caremani et al. [11,44] (Fig. 2 D) and the branched model of Debold and co-workers [27,30] (Fig. 2 E). For the multistep Pi-release model in Fig. 2C, the Pi-release indicated by yellow highlight, refers to Pi-release from secondary and tertiary sites. In all other models in Fig. 2, the Pi in yellow highlight indicates Pi-release from the active site. Below, we focus on the three alternative models as the conventional models have been researched in appreciable detail previously [2,7,9,10,12,13,16,20,28,52,53].

### Alternative model 1: Multistep orthophosphate release

The model with multistep Pi-release emerged from structural and reverse genetics studies of Llinas et al. [16] applied primarily to myosin VI. The central result in that study was evidence for the so-called Pi-release state, the state from which Pi leaves myosin [16]. This state, that had not been previously observed, is characterized by stereo-specific weak, actin-attachment that differs from the initial actin-attached state by having the nucleotide interacting switch-II in an open position. We later incorporated such a Pi-release state into a mechanokinetic model for myosin II operation [19,20], allowing us to decipher the mechanism of action of the small molecular myosin II inhibitor blebbistatin [20]. Moreover, structural modelling (Fig. 3A) corroborated the Pi-release properties of a similar state in myosin II [34]. Of further importance for the development of the multistep Pi-release model [34], Llinas et al. [16] found evidence for a secondary Pi binding sites outside the active site in myosin VI. We [34] used molecular modelling (Fig. 3B) to demonstrate a similar secondary Pi-binding site in myosin II. In addition we found evidence for a set of tertiary Pi-binding sites possibly related to a positive electrostatic surface potential on myosin (Fig. 3C). The latter evidence is based on single molecule studies showing that increased [Pi] competitively inhibited non-specific binding of fluorescent ATP to myosin [34]. The tertiary sites were not explicitly considered by Llinas et al. [16], but their existence in myosin II gained strong support from the single molecule competition assay. These results suggest overlap with non-specific ATP binding sites and positively charged residues (Fig. 3C-D). Altogether, there is strong evidence [2,16,54,55] that Pi, after its release from the active site, is guided via the so-called back door (Fig. 3A) to bulk solution while temporarily pausing in secondary and tertiary binding sites. The explanatory power of this multistep Pi-release model is substantial, accounting for a range of contractile phenomena that appear contradictory when interpreted in conventional models [34]. On the one hand, this includes the [Pi] independence of maximum sliding velocity and the monotonous decrease in isometric force with increased [Pi]. On the other hand, the model also accounts for the slow Pi-release [7,41] [16,42] and the simultaneous lack of effects of altered [Pi] on mechanical transients of muscle and isolated actomyosin [6,10,27]. Other strengths of the multistep Pi-release model [34] include available evidence for existence of both secondary and tertiary Pi-binding sites [34]. A limitation is that it is unclear if the kinetics of Pi-binding and subsequent release at secondary and tertiary sites is sufficiently slow to account for the delay (5-100 ms; varying between myosin isoforms) [7,16,41,42] of Pi-appearance in solution. It would also be of appreciable value to support the theoretical evidence for existence of the secondary Pi-binding site in myosin with experimental data. Moreover, it is not clear if the multistep Pi-release model accounts for changes in gliding velocity with increased [Pi] at low pH as suggested for the branched model (see below).

**Fig. 3.**
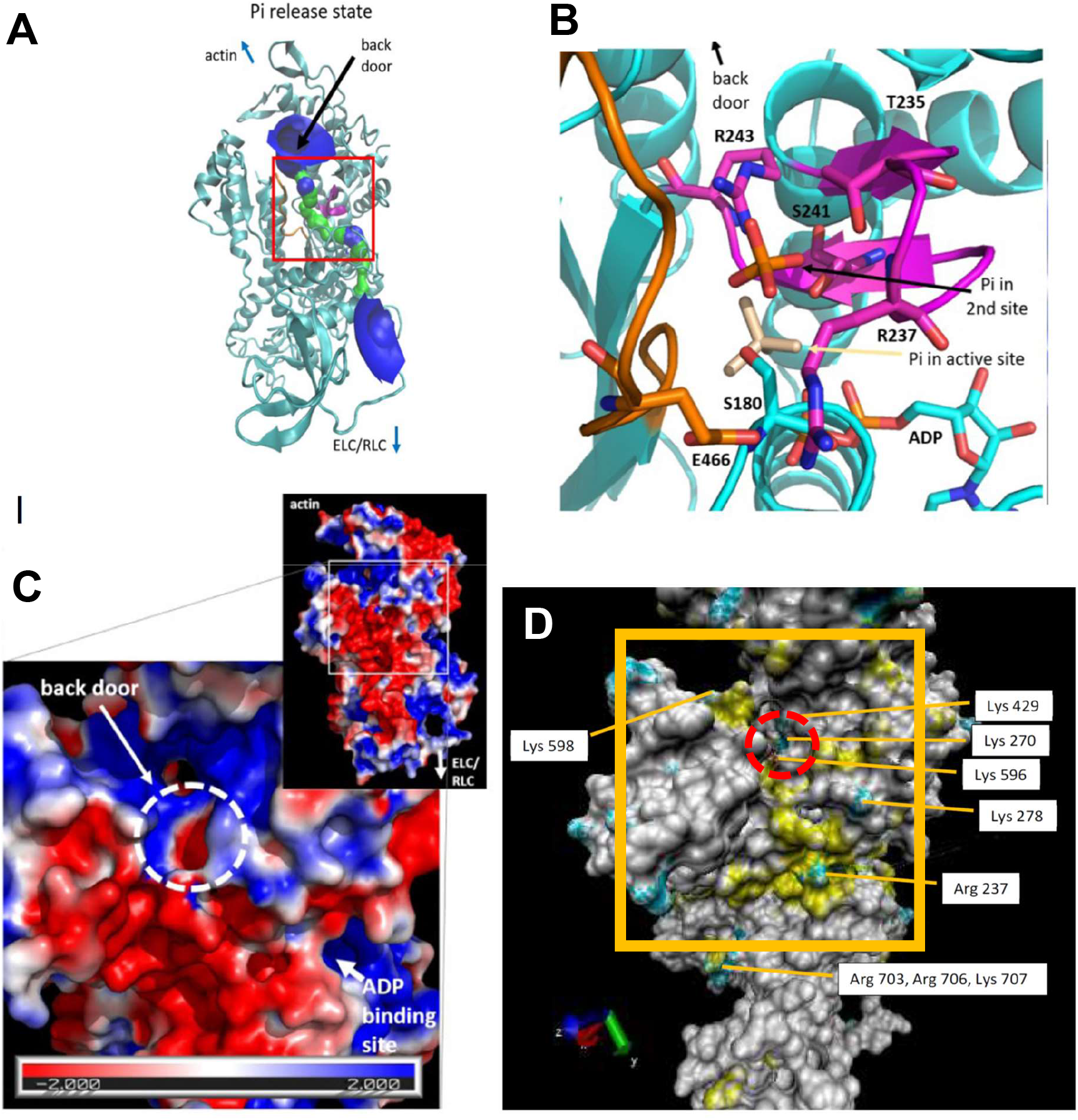
Molecular models of Pi-release path in cardiac β-myosin II. **A.** Model of the Pi-release state (from [34]) in bovine cardiac β-myosin II. The model is based on the bovine cardiac myosin II pre-power-stroke structure (PDB 5N6A) using the switch-II conformation from the Pi-release state of myosin VI (PDB 4PFO). The model depicts the Pi-release path via the “back door” in green and blue as identified using the program Hole (v. 2.2.005; http://www.holeprogram.org/) [56,57] Red box: Active site area illustrated in detail in **B.** Structural model of active site of myosin II in the Pi-release state modeled in A with a phosphate in the secondary site (orange) as well as in the active site (beige). To allow release of the phosphate, Switch-II (orange wire) has moved from its initial pre-powerstroke state to the modelled Pi-release state shown. Switch-I shown in purple. **C.** Electrostatics around the opening of the back door in myosin II (PDB: 5N6A [http://doi.org/10.2210/pdb5N6A/pdb] (Pre-powerstroke)). The surface is coloured according to its electrostatic potential, from -2.0 k_B_T (deep red) to +2.0 k_B_T (deep blue) indicating appreciable positive charge with the expected potential to bind Pi. **D.** The myosin II surface structure similar to that in C (PDB: 5N6A) but showing positively charged amino acids (blue) and ATP-binding areas (yellow) identified using the program ATPint [58] (for further details see [59]). Positively charged amino acids that overlap or are very close to identified ATP binding regions are labeled in boxes. Dashed circle indicates opening of back door as in C. Panels A-C reproduced from [34] under license CC BY 4.0. Panel D rendered using VMD (v. 1.9.3; http://www.ks.uiuc.edu/Research/vmd/ [60].

### Alternative model 2: The loose coupling model

The loose coupling model of Caremani et al. [11,44] was developed to account for the effects of varied [Pi] on transient and steady-state effects of changes in load on isometrically contracting muscle fibers. The general principles of the loose coupling model are illustrated in Fig. 2D with details [11,44] in Fig. 4. Its key features (Fig. 4) are: i. the power-stroke has three sub-strokes corresponding to one pre-power-stroke state (upper row in Fig. 4A) and three post-power-stroke states (M_2_-M_4_ in Fig. 4A) with strain-dependent transition rates in between, ii. each sub-stroke occurs with the same rate whether both ADP and Pi or only ADP is at the active site, iii. the release of Pi (step 4 in Fig. 4A) occurs before release of ADP (step 5 in Fig. 4A) but these processes are orthogonal to the power-strokes, i.e. Pi and ADP can be released from the active site in any of the pre-power-stroke (M_1_), or intermediate-final post-power-stroke states (M_2_-M_4_), iv. the Pi-release rate increases from <100 s^-1^ to 5000 s^-1^ under progression from the pre-power-stroke state over the two intermediate post-power stroke states to the final post-stroke state, v. the myosin motor can slip between two consecutive binding sites along the actin filament while it is at an intermediate state of its biochemical and structural (force-generating) cycle (Fig. 4B and steps 8 – 9 in Fig. 4A) and vi. myosin may undergo unconventional detachment from actin with the hydrolysis products still at the active site (step 7 in Fig. 4A), followed by a fast release of the products and binding of a new ATP. The latter feature was originally introduced [61], to explain a smaller reduction with increased [Pi] in ATP turnover rate than in tension development during isometric contraction [33,62,63].

**Fig. 4.**
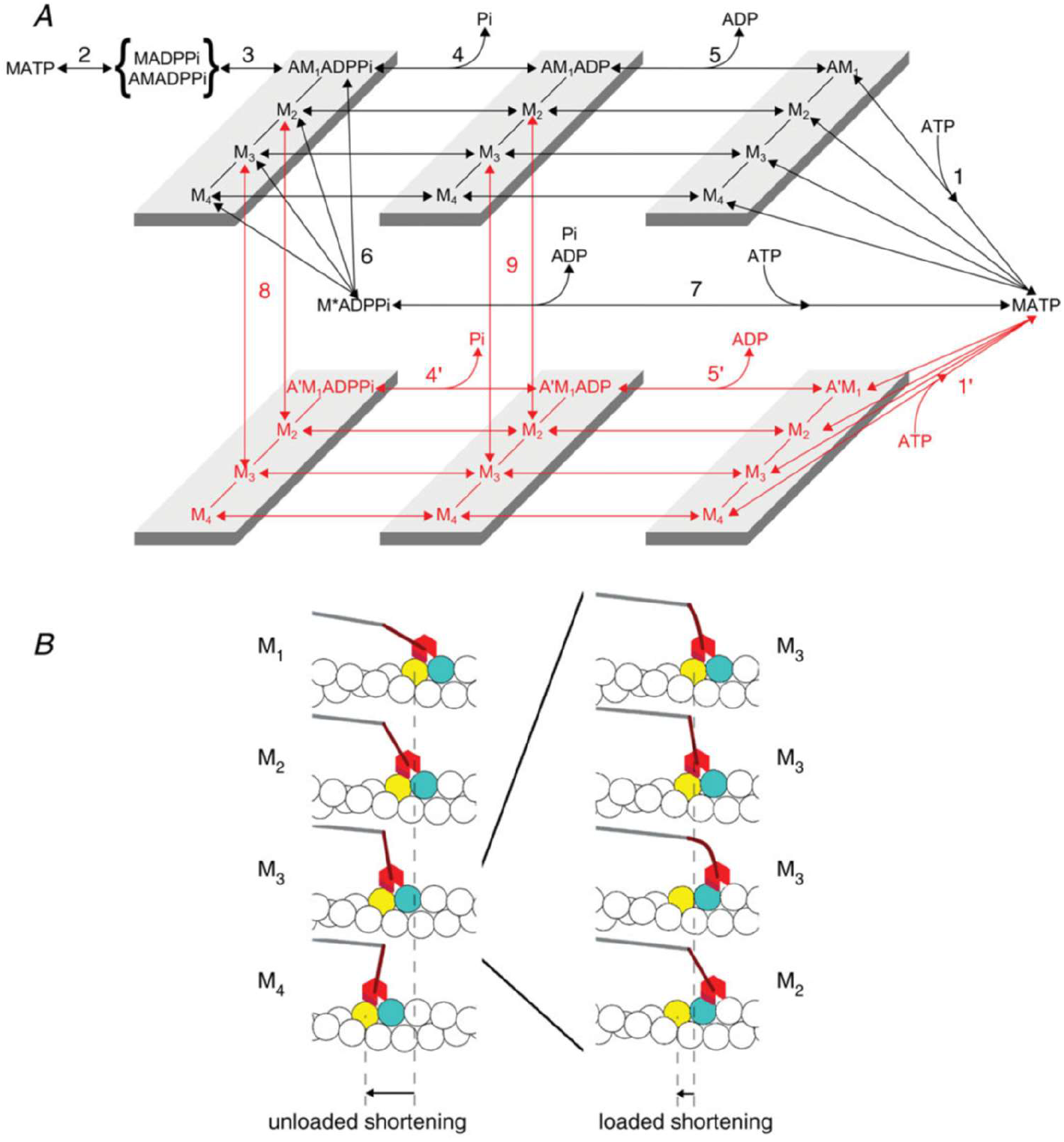
Details of the cross-bridge cycle in the loose coupling model of Caremani et al. **A.** Model states with five different myosin (M) and actomyosin (AM) nucleotide (N) states with either ATP (far left), ADP and Pi, ADP or no nucleotide (far right; “rigor-state”). In the attached (AMN) states there are four different structural states (M_1_-M_4_; see further panel B) each with the lever arm for each nucleotide state with the lever arm in different positions corresponding to three sub-strokes of the power-stroke. Additionally, slippage transitions may occur (transitions 8 and 9) from a first attachment position along the actin filament (top, black; left in panel B) to at least one other position along actin (bottom, red; right in panel B). Finally, it is also assumed that unconventional detachment (transition 6) may occur. **B.** Structural states (M_1_-M_4_) existing for each AMN state. Binding to only one actin site (left; black in A) possible during isometric contraction but slippage to neighboring sites (right; red in A) is assumed to occur during loaded shortening. The figure is reproduced from [11] with permission from WILEY.

The loose coupling model differs from the multistep Pi-release model by its assumption of three sub-strokes, instead of just one, and slippage of the myosin head between neighboring actin filament sites. These features are, however, not relevant in relation to the Pi-release mechanism and could in principle be introduced also in the multi-step Pi-release model. The most important differences between the loose coupling model and the multistep Pi-release model in relation to Pi-release and the power-stroke(s) is how the orthogonal biochemical (Pi and ADP release) and mechanical (power-stroke) cycles are implemented. In the loose-coupling model, the partial uncoupling between the two cycles is assumed to occur without the assumption of secondary and tertiary Pi-binding sites. In contrast, the latter sites are central in the multistep Pi-release model [34]. With lack of the secondary and tertiary sites in the loose coupling model, the Pi-release from the active site must be slow in order to account for the slow appearance of Pi in solution [7,16,41,42] as this effect cannot be attributed to binding outside the active site. Moreover, contrary to the predictions of Houdusse, Sweeney and co-workers [2,16], Pi release from the active site cannot be a necessary condition for occurrence of the sub-strokes in the loose coupling model. Despite these differences, however, the loose-coupling model seems to give similar predictions as the multistep Pi-release model [34] in several respects. This includes [Pi] independence of both the power-stroke rate and the maximum velocity of shortening [11]. Presumably, the model also predicts a monotonous reduction in isometric force with increased [Pi] although this was not explicitly tested for a range of Pi-concentrations. However, it is not clear that the model can explain the findings that Pi-release for actomyosin in solution is appreciably slower than the power stroke [7]. In the loose-coupling model, the Pi-release rate is taken as > 1000 s^-1^ for states towards the end of the power-stroke (presumably to account for the high shortening velocity). These states (e.g. M_4_ in Fig. 4A) are likely to be heavily populated in solution experiments where there are no elastic strains that prevent the power stroke. Therefore, one would expect the loose-coupling model [11] to predict a negligible delay between the power stroke and the Pi-release, contrary to what was observed by Muretta et al. [7].

To summarize, a challenge related to the loose coupling model is whether it can simultaneously account for the low Pi-release rate observed in biochemical studies and the fast velocity of shortening observed in muscle and in vitro motility assays. A question that also must be answered is if the power-stroke can occur with Pi in the active site.

### Alternative model 3: Branched model

The branched model of Debold, Walcott and others [18,27,30,31] is characterized by: i. a branched kinetic pathway with Pi-binding to a post-power-stroke actin-attached cross-bridge state, leading to detachment of the myosin head into a state with the lever arm in the post-power-stroke position (Fig. 2E). ii. Pi binding to the active site in the rigor (AM) state, competing with ATP binding to the active site. In a recent version of the model [27], initial Pi-release is believed to occur after the power-stroke (cf. Fig. 2E) which seems logical considering that Pi-re-binding is assumed to occur in the post-power-stroke state.

The main discriminating feature between this model, the multistep Pi-release model [34] and most other models, is the branched pathway (i). The other discriminating feature, the inhibition by Pi of ATP binding to the active site in the rigor state (ii), is not unique to this model (see e.g. [34,64]); we will not consider it further as it is not directly relevant for the Pi-release mechanism associated with the power-stroke. Regarding the branching, a similar pathway (with somewhat different properties) was considered for isometric contraction by Linari et al. [61] using a simple kinetic scheme. In the more complete mechanokinetic analysis of that group [11,44], giving the loose coupling model, the branched pathway only plays a secondary role. We therefore focus on the version put forward by Debold, Walcott and co-workers [18,27,30,31].

The main motivations behind the branched model were to account for two findings that are difficult to explain using conventional models [18]: First, that increased [Pi] reduces isometric force development appreciably more than the ATP turnover rate during isometric contraction and second, that the maximum sliding velocity in the in vitro motility assay is increased with increased [Pi] at low pH (<7). With regard to the first of these findings, a serial kinetic scheme, without consideration of cross-bridge elasticity (e.g., no x-dependence of rate constants in Fig. 2A-B) was found to predict [61] a larger reduction of the isometric ATP turnover rate than of isometric force upon increased [Pi]. This effect is in striking contrast to what is seen experimentally with small reduction in the isometric ATP turnover rate compared to isometric force with increased [Pi]. This discrepancy could be accounted for by introducing a branched pathway as in Fig. 2E [61].

The second motivation for the branched model was that the gliding velocity in the in vitro motility assay increases in response to increased [Pi] at low pH. The authors [30] proposed that Pi-binding to the post-power-stroke AMADP state, followed by detachment into a post-power-stroke state, would explain this result. They relied on findings [65] that the AMADP state is prolonged compared to the AM (rigor) state at low pH and high [ATP] making the lifetime of the AMADP state more important in limiting the detachment rate and thereby the maximum actin gliding velocity. This would explain that Pi-induced detachment from the AMADP state has greater effect on the velocity [30] at low pH. Subsequent mechanokinetic versions of the branched model seem to also account for a range of additional effects of varied [Pi] [31].

However, the branched model also has some limitations. First, the lower effect of varied [Pi] on the isometric ATP turnover rate than on isometric force is not unique to models with branched pathways. The effect is an inherent component of a range of other mechanokinetic models [19,28,38] that explicitly include effects of the cross-bridge elasticity (e.g. strain-dependence of rate constants). As outlined above, the strain dependence can be described by a variable x. If this variable is defined as giving the strain of the elastic element in the post-power-stroke cross-bridge state, the strain in the pre-power-stroke state is equal to x-h if h is the size of the power-stroke and if there is only one power-stroke (cf. [49]). Assuming such a simple model, cross-bridge attachment into the pre-power-stroke state occurs primarily for large x (around x=h) whereas, cross-bridge detachment (including ADP release and ATP re-binding) from the post-power-stroke state primarily occurs at low x. The cross-bridges that carry highest force are those in the post-power-stroke state at high x, together with cross-bridges in their pre-power-stroke state with high strain already at attachment. These high-force cross-bridges are, however, least prone to undergo detachment by ADP release and ATP binding (due to strain-dependence of rate constants) but most prone to undergo detachment by reversal of the power-stroke as a result of Pi-rebinding (cf. [19,66,67]). In isometric contraction, this predicts that increased [Pi] alters the steady-state cross-bridge distribution as illustrated in Fig. 5 for the model in [19] (similar to Fig. 2A). Only cross-bridges at high x-values (at x ≈ h and higher) are lost with increased [Pi]. These cross-bridges would not readily detach via ADP release and ATP re-binding. Thus, increased [Pi] causes a large drop in isometric force but, in this case, a negligible change in ATP turnover rate (limited by ADP release under isometric conditions), in general agreement with experimental results. Similar predictions that much of the isometric force is held by pre-power-stroke cross-bridges follows from theoretical arguments [68].

**Fig. 5.**
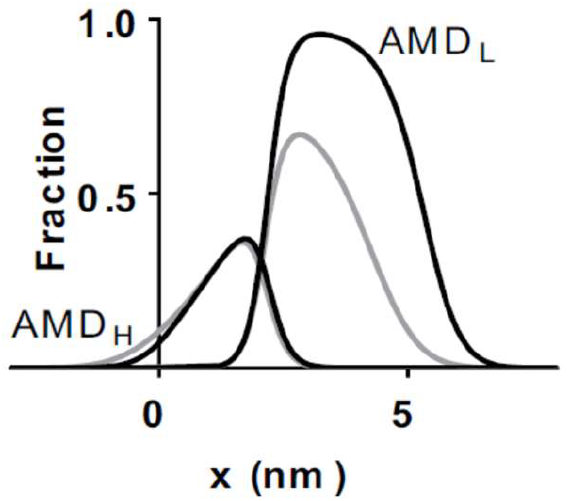
Population of pre-power-stroke (low-force; AMD_L_) and post-power-stroke (high-force; AMD_H_) states as function of cross-bridge strain parameter x in isometric contraction. This figure corresponds to the kinetic scheme in Fig. 2A. Strain of the elastic element in the AMD_L_ state is 6.7 nm lower (corresponding to power-stroke size, h=6.7 nm) than in the AMD_H_ state. Pi can rebind to the AMD_L_ state (but not to the AMD_H_ state) in this model, leading to reversal of the cross-bridge attachment etc. In contrast, ADP can be released only from the AMD_H_ state following a strain dependent transition. The fraction of attached cross-bridges relates to all cross-bridges (detached and attached) for each given value of x. Modified from [19] under license CC BY 4.0.

Regarding the [Pi]-effects on velocity at low pH, there are also some limitations that need to be considered. First, to the best of our knowledge, the increased velocity in response to increased [Pi] at low pH has only been demonstrated experimentally using in vitro experiments on isolated proteins [30]. These results may differ from those obtained in muscle fiber experiments (cf. [50,69–71]). Accordingly, the maximum shortening velocity in muscle cells, in contrast to the in vitro motility assay data in [30], seem to be little affected by altered [Pi] at low pH [17,72]. Indeed, some studies have even found evidence that increased [Pi] and lowered pH are synergistic in reducing velocity in muscle fibers [73]. These issues need to be further elucidated.

Even if the experimental [Pi]-pH velocity relationship is consistent with the branched kinetic model, Sleep and Hutton (15) and later Bowater and Sleep (16) found that Pi-rebinding to myosin II leads to re-synthesis of ATP. This is consistent with a reversal of the attachment step and the ATP hydrolysis step on the myosin active site (as in all models in Fig. 2 other than the branched model). The mechanism with Pi-induced detachment into a post-power-stroke state is difficult to reconcile with these findings.

Effects of an S217A mutation in myosin V (S236A in myosin II and S203A in myosin VI) have been taken as support for the branched model, combined with initial Pi release after the power-stroke. Forgacs et al. [74] found an appreciable slowing of the Pi-release step in myosin V with an S217A mutation. This finding was similar to that of Llinas et al. [16] for the corresponding mutation in myosin VI and myosin II. Scott et al. [27] used optical trapping to study the effects of the myosin V S217A mutation on the power-stroke in single molecules. They found that, both in the presence and absence of 30 mM Pi, the wild-type myosin V produced a fast power-stroke (≥ 500 s^-1^) without detectable delay following actin binding, but with the 25 % longest binding events eliminated by added Pi. Further, they found that the S217A myosin V construct generated a power-stroke following binding to actin that was similar in rate and size as for wild-type myosin V. All these findings were taken as evidence [27] that myosin V generates its power-stroke with Pi in its active site and that Pi-re-binding leads to detachment into a post-power-stroke state as proposed in the branched model. However, these results warrant interpretation within the framework of different models. First, even if the S217A mutation prevents (or slows) the entrance from the active site into the back door, leading to slower Pi-release, it is not clear if the Pi would stay in the active site for longer time due to this effect. Instead, Pi may be trapped in an intermediate position like the secondary or tertiary sites in the multistep Pi-release model [34]. Indeed, the corresponding serines in myosin VI [16] and myosin II [34] do not only communicate with the active site but also participate in Pi-coordination in the secondary Pi-binding site outside the active site. Accordingly, the effect of the S217A mutation on Pi-release did not limit Houdusse, Sweeney and co-workers to conclude that the Pi must leave the active site before the power-stroke [2,16].

### Testing the alternative models for Pi-release

#### Multistep Pi-release model

Despite evidence for secondary and tertiary Pi-binding sites, it is not clear if the Pi-binding exhibits sufficiently slow kinetics to account for the observed slow rate of Pi-release [7,16,41,42]. If the off rate from the secondary and tertiary sites is faster than the overall Pi-release rate, it would suggest that the rate limitation of Pi-release is the dissociation from the active site, arguing against the multistep Pi-release model. We proposed [34] that this may be tested by mutation of the secondary and tertiary sites in ways that either change the kinetics of the binding or prevent Pi-binding to the sites. We also proposed that the double mutation R243E/E466R in cardiac myosin II could be useful in this regard [75] and preferred over the single R243E mutation (disrupting the secondary Pi-site). This is because the double mutation is expected to preserve other aspects of actomyosin function [75].

We have found that both the R243E and R243E/E466R mutations are properly transfected into C2C12 cells by a new virus free method [76] that would allow convenient screening of a range of mutants. Both mutants also express well in these cells (Fig. 6). We will next perform basic functional characterization and quantification of the Pi-release rate constant. The advantage of using cardiac myosin II for these studies compared to Dictyostelium myosin is that the R243 residue is also a hotspot for disease causing mutations, e.g. in cardiomyopathies [77,78].

**Fig. 6.**
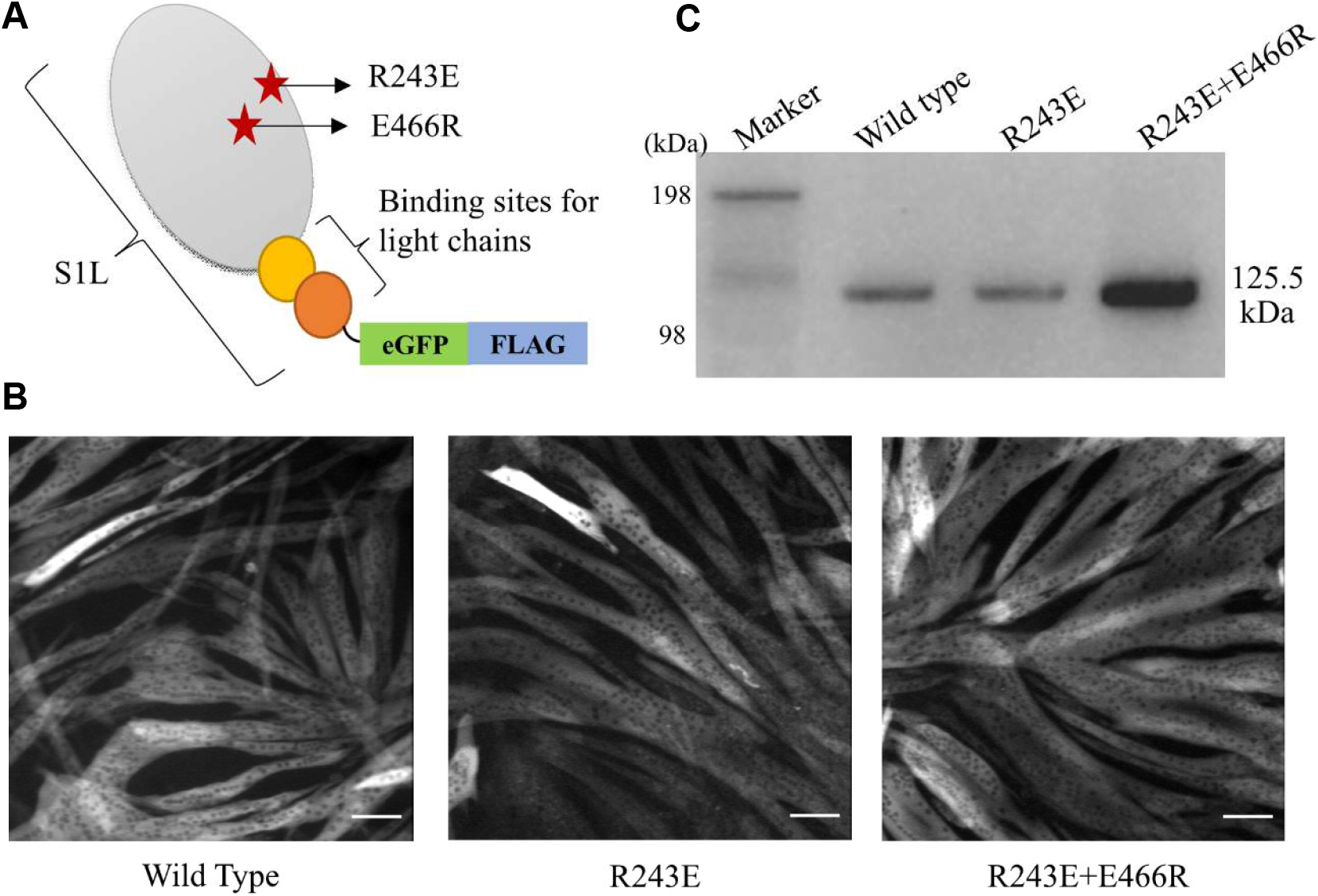
Virus-free transfection, expression, and purification of human β-cardiac myosin constructs in C2C12 cells. **A**. Cartoon of expression construct representing β-cardiac myosin II motor domain containing a single myosin heavy chain (MHC) fragment (S1L) with binding sites for essential and regulatory light chains fused with eGFP and a FLAG tag. Stars indicate mutations R243E and E466R. **B**. EGFP fluorescence observed in differentiated C2C12 myotubes, expressing wild type, R243E and R243E+E466R mutant proteins. Bar, 100 µm. **C.** Western blot analysis of wild-type and mutant S1L-eGFP-FLAG protein (125.5 kDa) purified from single 60 mm culture dish per construct and detected via anti FLAG antibodies.

The multistep Pi-release model predicts that secondary Pi-binding accounts for a major fraction of a delay of up to 100 ms before Pi appears in solution after the power-stroke (data from bovine cardiac β-myosin [43]). Therefore, we expect that the R243E/E466R mutation, disrupting the secondary site, would appreciably increase the Pi-dissociation rate from 10 s^-1^ or somewhat higher in wild-type human β-myosin to possibly 1000 s^-1^.

Recently, [34] we raised the possibility that also Pi-binding to the tertiary sites may contribute significantly to the observed delay of Pi-appearance in solution. This should be testable using a similar strategy as for the secondary site, i.e., quantification of the Pi-release rate of β-myosin constructs with tertiary site mutations designed to prevent Pi-binding. However, the location of the tertiary sites is less well-defined. On the one hand, we noted a positive surface electrostatic potential around the opening of the back door for Pi release (Fig. 3C). On the other hand, we found that the Pi binding outcompeted almost all non-specific Alexa647-ATP binding to the surface of the myosin head. It is not clear if these Alexa647-ATP binding sites include the positively charged regions around the back door opening. This possibility is analyzed for bovine cardiac β-myosin in Fig. 3D. Here, we indicate all positively charged amino acids (blue) on the myosin surface close to the back door opening together with residues (yellow) that are involved in non-specific ATP binding according to the program ATPint ([58]; see also [59]). Specifically, we label positively charged amino acids in Fig. 3D that are parts of the non-specific ATP-binding sites or very close to these. Such positively charged amino acids would be interesting candidates for mutations to disrupt tertiary Pi-binding sites. Importantly, the positive charged residues labelled in Fig. 3D are all conserved between bovine (Fig. 3) and human cardiac β-myosin (sequence alignment in ClustalW).

After mutation of secondary and putative tertiary Pi-binding sites, according to the above strategy, and further strategies used in similar studies of another molecule [79], we will first characterize the general functionality. After initial characterization of mutants we will select those with maintained overall function of greatest interest for further detailed studies of the Pi-release.

#### Loose coupling model

Testable predictions of the loose coupling model include: i. Pi must be allowed to leave the active site either before or after the power-stroke, ii. the secondary and tertiary Pi-binding sites play no role in delaying the Pi-appearance in solution and iii. the rates of Pi-release from the pre- and the three post-power-stroke states must be such that both the maximum velocity of shortening and the delay of Pi-appearance in solution after the power-stroke are accounted for. The first of these predictions (i) has been partly tested in high-speed AFM recordings [34] and single molecule force measurements [39], suggesting that the power-stroke can occur without Pi in the active site. What remains to be shown is that the reverse is also true, i.e., that the power-stroke can occur with Pi in the active site. If Pi release from the active site is very fast, it may be challenging to test the latter possibility. However, the loose coupling model assumes rather slow Pi-release from the active site suggesting that it should be in principle possible to detect Pi binding both before and after the power-stroke (at least after the first sub-strokes). It is presently not clear how it would be possible to monitor such binding but changes in internal protein fluorescence upon Pi-release from the active site may be an option [22]. Testing the second prediction (ii) that the secondary and tertiary sites do not contribute to delayed appearance of Pi in solution was considered above. Thus, if mutations of the secondary and tertiary sites do not modify the delayed Pi-appearance in bulk solution, the remaining possibility seems to be that the delay is due to slow dissociation from the active site as in the loose coupling model. However, in relation to the third testable prediction (iii), it then also remains to be shown that both such a slow Pi-appearance and a high maximum shortening velocity can be explained by the same set of model parameter values. Thus, the mechanokinetic model (as in [11]) with this given set of parameter values should account not only for the maximum velocity of shortening at varied [Pi] but also for the slow appearance of Pi in solution after the power-stroke as seen in [7].

#### Branched model

Most importantly, the findings of Sleep and Hutton [32] and later, Bowater and Sleep [33] that ATP is resynthesized upon Pi-binding need to be explained within the framework of the branched model. These results of Sleep and co-workers, seem to suggest that Pi-rebinding leads to reversal of the original Pi-release and attachment steps, rather than to detachment via a branched pathway. In such models (those in Figs. 2 except the branched model in Fig. 2E), one would expect a nucleotide dwell time distribution in single molecule studies [59,64,80] with more slow events upon increased [Pi]. This is because some of the cross-bridges that are detached through reversal of the Pi-release and the attachment steps may eventually go through several back-and-forth transitions with reversals upon Pi-binding followed by reattachment of myosin to actin, new Pi-induced reversals etc. Indeed, the final release of Pi from myosin may occur from detached heads, a process that is 100-fold slower than actin-activated Pi-release [21]. Such an effect would be expected to be most prominent in conditions with already prolonged dwell-times in low-force states (e.g. AMD_L_ and AMDP_L_ in Fig. 2). These are particularly highly populated under isometric conditions. One would therefore expect most substantial increase in the number of slow events upon increased [Pi] under such conditions. Similar effects of increased [Pi] are not expected in the branched pathway if the Pi-re-binding and subsequent product release rates (lower row of Fig. 2E) are as fast as assumed in a recent version of the model [31] (see also [61]). Evidence suggesting more slow ATP dwell-times in an isometric single molecule assay upon increased [Pi], was provided by Amrute-Nayak et al. ([64]). A recently modified version [59] of their isometric single molecule ATPase assay is likely better at detecting long events due to removal of artifacts that tend to bias the dwell-times towards short events. Therefore, in order to better distinguish between branched and unbranched models, it would be of interest to repeat the previous experiments [64] using methods in [59] to more definitely corroborate the idea that increased [Pi] increases the fraction of long fluorescent nucleotide dwell-times. It is important to note, in this connection, that both the branched model and the multistep Pi-release model would account for loss of the longest actin-binding events upon increased [Pi] in optical tweezers studies of myosin V [27]. Thus, the longest actin-binding events may be equally likely eliminated by detachment into a post-power-stroke state and by reversal of the cross-bridge attachment. Only in the latter case, however, would the ATP dwell-times be expected to increase as explained above.

#### General tests

A further way to evaluate effects of Pi-release and binding events would be to monitor effects of the release of caged Pi from wild type and mutated myosin in real-time using either optical tweezers or hs-AFM. Such methods simultaneously capture the structural and dynamic features of the myosin molecule bound to actin, allowing tests of various models of myosin II activity in the different experimental conditions. Whereas optical trapping methods would have sufficient time resolution to capture relevant mechanical events following Pi-release, the hs-AFM method is currently limited in temporal resolution, preventing visualization of the fast sub-steps during power stoke and Pi release. It has the potential to study structural changes related to Pi-binding; we hope to soon bypass time limitations by applying a single-line scanning method. This approach enables image scanning, while the AFM cantilever probe is at a fixed *x–y* position. It allows monitoring of the height fluctuations under the cantilever probe in the *z*-direction, producing single-line height data (*x, z, t*) with Angstrom spatial and microsecond (μs) temporal resolution [81].

## Conclusions

We describe three alternative models for Pi-release with either a branched pathway or partial uncoupling between the biochemical cycle and the power-stroke so that the Pi may be released from myosin either before or after the power-stroke. The latter two “uncoupled” models are distinguished from each other primarily by the importance of secondary and tertiary Pi-binding sites to account for delayed Pi-appearance in solution. These sites are central in the multistep Pi-release model but play no roles in the loose coupling model.

Considering the current uncertainties it is essential to apply critical tests to predictions of the alternative models. In depth insight into the Pi-release mechanism is important not only for full insight into the fundamental mechanisms of energy transduction by actin and myosin. It is also increasingly important because this mechanism is targeted by several small molecular myosin active compounds with therapeutic potential [43,82–85] in cancer [86], heart failure[87], cardiomyopathies [88,89], and skeletal muscle disorders [90].

## Acknowledgements

This work was funded by The Swedish Research Council (grant number 2019-03456), The Faculty of Health and Life Sciences at The Linnaeus University, Sweden.

## Data availability

The data that support the findings of this study are available from the corresponding author upon reasonable request.

## References

1 Heissler, SM, Sellers, JR. 2016. Kinetic Adaptations of Myosins for Their Diverse Cellular Functions. Traffic 17: 839–59.

2 Robert-Paganin, J, Pylypenko, O, Kikuti, C, Sweeney, HL, et al. 2020. Force Generation by Myosin Motors: A Structural Perspective. Chem Rev 120: 5–35.

3 Trivedi, DV, Nag, S, Spudich, A, Ruppel, KM, et al. 2020. The Myosin Family of Mechanoenzymes: From Mechanisms to Therapeutic Approaches. Annu Rev Biochem 89: 667–93.

4 Zhao, Y, Kawai, M. 1994. Kinetic and thermodynamic studies of the cross-bridge cycle in rabbit psoas muscle fibers. Biophys J 67: 1655–68.

5 Coupland, ME, Puchert, E, Ranatunga, KW. 2001. Temperature dependence of active tension in mammalian (rabbit psoas) muscle fibres: effect of inorganic phosphate. J Physiol 536: 879–91.

6 Ranatunga, KW, Coupland, ME, Mutungi, G. 2002. An asymmetry in the phosphate dependence of tension transients induced by length perturbation in mammalian (rabbit psoas) muscle fibres. J Physiol 542: 899–910.

7 Muretta, JM, Rohde, JA, Johnsrud, DO, Cornea, S, et al. 2015. Direct real-time detection of the structural and biochemical events in the myosin power stroke. Proc Natl Acad Sci U S A 112: 14272–7.

8 Trivedi, DV, Muretta, JM, Swenson, AM, Davis, JP, et al. 2015. Direct measurements of the coordination of lever arm swing and the catalytic cycle in myosin V. Proc Natl Acad Sci U S A 112: 14593–8.

9 Dantzig, JA, Goldman, YE, Millar, NC, Lacktis, J, et al. 1992. Reversal of the cross-bridge force-generating transition by photogeneration of phosphate in rabbit psoas muscle fibres. J Physiol 451: 247–78.

10 Woody, MS, Winkelmann, DA, Capitanio, M, Ostap, EM, et al. 2019. Single molecule mechanics resolves the earliest events in force generation by cardiac myosin. Elife 8.

11 Caremani, M, Melli, L, Dolfi, M, Lombardi, V, et al. 2013. The working stroke of the myosin II motor in muscle is not tightly coupled to release of orthophosphate from its active site. J Physiol 591: 5187–205.

12 Governali, S, Caremani, M, Gallart, C, Pertici, I, et al. 2020. Orthophosphate increases the efficiency of slow muscle-myosin isoform in the presence of omecamtiv mecarbil. Nature communications 11: 3405.

13 Tesi, C, Colomo, F, Nencini, S, Piroddi, N, et al. 2000. The effect of inorganic phosphate on force generation in single myofibrils from rabbit skeletal muscle. Biophys J 78: 3081–92.

14 Tesi, C, Colomo, F, Piroddi, N, Poggesi C. 2002. Characterization of the cross-bridge force-generating step using inorganic phosphate and BDM in myofibrils from rabbit skeletal muscles. J Physiol 541: 187–99.

15 Stehle R. 2017. Force Responses and Sarcomere Dynamics of Cardiac Myofibrils Induced by Rapid Changes in [Pi]. Biophys J 112: 356–67.

16 Llinas, P, Isabet, T, Song, L, Ropars, V, et al. 2015. How actin initiates the motor activity of Myosin. Dev Cell 33: 401–12.

17 Karatzaferi, C, Adamek, N, Geeves MA. 2017. Modulators of actin-myosin dissociation: basis for muscle type functional differences during fatigue. American journal of physiology Cell physiology 313: C644–C54.

18 Debold, EP, Walcott, S, Woodward, M, Turner, MA. 2013. Direct observation of phosphate inhibiting the force-generating capacity of a miniensemble of Myosin molecules. Biophys J 105: 2374–84.

19 Månsson A. 2021. The effects of inorganic phosphate on muscle force development and energetics: challenges in modelling related to experimental uncertainties. J Muscle Res Cell Motil 44: 33–46.

20 Rahman, MA, Usaj, M, Rassier, DE, Månsson A. 2018. Blebbistatin Effects Expose Hidden Secrets in the Force-Generating Cycle of Actin and Myosin. Biophys J 115: 386–97.

21 Malnasi-Csizmadia, A, Kovacs M. 2010. Emerging complex pathways of the actomyosin powerstroke. Trends Biochem Sci 35: 684–90.

22. Gyimesi M, Kintses B, Bodor A, Perczel A, et al. 2008. The mechanism of the reverse recovery, step, phosphate, release, and actin activation of Dictyostelium myosin II. J Biol Chem 283: 8153–63.

23 Cooke, R, Pate E. 1985. The effects of ADP and phosphate on the contraction of muscle fibers. Biophys J 48: 789–98.

24 Månsson, A, Rassier, D, Tsiavaliaris G. 2015. Poorly Understood Aspects of Striated Muscle Contraction. BioMed Research International 2015: 28.

25 Gunther, LK, Rohde, JA, Tang, W, Cirilo, JA, Jr., et al. 2020. FRET and optical trapping reveal mechanisms of actin activation of the power stroke and phosphate release in myosin V. J Biol Chem 295: 17383–97.

26 Stehle, R, Tesi C. 2017. Kinetic coupling of phosphate, release, force generation and rate-limiting steps in the cross-bridge cycle. J Muscle Res Cell Motil.

27 Scott, B, Marang, C, Woodward, M, Debold EP. 2021. Myosin’s powerstroke occurs prior to the release of phosphate from the active site. Cytoskeleton (Hoboken).

28 Smith DA. 2014. A new mechanokinetic model for muscle, contraction, where force and movement are triggered by phosphate release. J Muscle Res Cell Motil 35: 295–306.

29 Homsher E. 2017. A New and Improved View of Force Production. Biophys J 112: 205–6.

30 Debold, EP, Turner, MA, Stout, JC, Walcott S. 2011. Phosphate enhances myosin-powered actin filament velocity under acidic conditions in a motility assay. Am J Physiol Regul Integr Comp Physiol 300: R1401–8.

31 Jarvis, K, Woodward, M, Debold, EP, Walcott S. 2018. Acidosis affects muscle contraction by slowing the rates myosin attaches to and detaches from actin. J Muscle Res Cell Motil 39: 135–47.

32 Sleep, JA, Hutton RL. 1980. Exchange between inorganic phosphate and adenosine 5’-triphosphate in the medium by actomyosin subfragment 1. Biochemistry (Mosc) 19: 1276–83.

33 Bowater, R, Sleep J. 1988. Demembranated muscle fibers catalyze a more rapid exchange between phosphate and adenosine triphosphate than actomyosin subfragment 1. Biochemistry 27: 5314–23.

34 Moretto, L, Usaj, M, Matusovsky, O, Rassier, DE, et al. 2022. Multistep orthophosphate release tunes actomyosin energy transduction. Nature communications 13: 4575.

35 Shalabi, N, Persson, M, Mansson, A, Vengallatore, S, et al. 2017. Sarcomere Stiffness during Stretching and Shortening of Rigor Skeletal Myofibrils. Biophys J 113: 2768–76.

36 Trivedi, DV, Adhikari, AS, Sarkar, SS, Ruppel, KM, et al. 2018. Hypertrophic cardiomyopathy and the myosin mesa: viewing an old disease in a new light. Biophysical reviews 10: 27–48.

37 Eisenberg, E, Hill, TL, Chen Y. 1980. Cross-bridge model of muscle contraction. Quantitative analysis. Biophys J 29: 195–227.

38 Pate, E, Cooke R. 1989. A model of crossbridge action: the effects of, ATP, ADP and Pi. J Muscle Res Cell Motil 10: 181–96.

39 Hwang, Y, Washio, T, Hisada, T, Higuchi, H, et al. 2021. A reverse stroke characterizes the force generation of cardiac, myofilaments, leading to an understanding of heart function. Proc Natl Acad Sci U S A 118.

40 Muretta, JM, Petersen, KJ, Thomas DD. 2013. Direct real-time detection of the actin-activated power stroke within the myosin catalytic domain. Proc Natl Acad Sci U S A 110: 7211–6.

41 Rohde, JA, Thomas, DD, Muretta JM. 2017. Heart failure drug changes the mechanoenzymology of the cardiac myosin powerstroke. Proc Natl Acad Sci U S A 114: E1796–E804.

42 White, HD, Belknap, B, Webb MR. 1997. Kinetics of nucleoside triphosphate cleavage and phosphate release steps by associated rabbit skeletal, actomyosin, measured using a novel fluorescent probe for phosphate. Biochemistry 36: 11828–36.

43 Swenson, AM, Tang, W, Blair, CA, Fetrow, CM, et al. 2017. Omecamtiv Mecarbil Enhances the Duty Ratio of Human beta-Cardiac Myosin Resulting in Increased Calcium Sensitivity and Slowed Force Development in Cardiac Muscle. J Biol Chem 292: 3768–78.

44 Caremani, M, Melli, L, Dolfi, M, Lombardi, V, et al. 2015. Force and number of myosin motors during muscle shortening and the coupling with the release of the ATP hydrolysis products. J Physiol 593: 3313–32.

45 Hill TL. 1974. Theoretical formalism for the sliding filament model of contraction of striated muscle. Part I. Prog Biophys Mol Biol 28: 267–340.

46 Eisenberg, E, Hill TL. 1978. A cross-bridge model of muscle contraction. Prog Biophys Mol Biol 33: 55–82.

47 Huxley AF. 1957. Muscle structure and theories of contraction. Prog Biophys Biophys Chem 7: 255–318.

48 Huxley, AF, Simmons RM. 1971. Proposed mechanism of force generation in striated muscle. Nature 233: 533–8.

49 Månsson, A, Usaj, M, Moretto, L, Rassier DE. 2018. Do Actomyosin Single-Molecule Mechanics Data Predict Mechanics of Contracting Muscle? International journal of molecular sciences 19.

50 Månsson A. 2020. Hypothesis: Single Actomyosin Properties Account for Ensemble Behavior in Active Muscle Shortening and Isometric Contraction. International journal of molecular sciences 21.

51 Eisenberg, E, Greene LE. 1980. The relation of muscle biochemistry to muscle physiology. Annu Rev Physiol 42: 293–309.

52 Månsson A. 2016. Actomyosin based contraction: one mechanokinetic model from single molecules to muscle? J Muscle Res Cell Motil 37: 181–94.

53 Offer, G, Ranatunga KW. 2020. The Location and Rate of the Phosphate Release Step in the Muscle Cross-Bridge Cycle. Biophys J 119: 1501–12.

54 Mugnai, ML, Thirumalai D. 2021. Step-Wise Hydration of Magnesium by Four Water Molecules Precedes Phosphate Release in a Myosin Motor. The journal of physical chemistry B 125: 1107–17.

55 Cecchini, M, Alexeev, Y, Karplus M. 2010. Pi release from myosin: a simulation analysis of possible pathways. Structure 18: 458–70.

56 Smart, OS, Goodfellow, JM, Wallace BA. 1993. The pore dimensions of gramicidin A. Biophys J 65: 2455–60.

57 Smart, OS, Neduvelil, JG, Wang, X, Wallace, BA, et al. 1996. HOLE: a program for the analysis of the pore dimensions of ion channel structural models. J Mol Graph 14: 354–60, 76.

58 Chauhan, JS, Mishra, NK, Raghava GP. 2009. Identification of ATP binding residues of a protein from its primary sequence. BMC Bioinformatics 10: 434.

59 Usaj, M, Moretto, L, Vemula, V, Salhotra, A, et al. 2021. Single molecule turnover of fluorescent ATP by myosin and actomyosin unveil elusive enzymatic mechanisms. Commun Biol 4: 64.

60 Humphrey, W, Dalke, A, Schulten K. 1996. VMD: visual molecular dynamics. J Mol Graph 14: 33-8, 27–8.

61 Linari, M, Caremani, M, Lombardi V. 2010. A kinetic model that explains the effect of inorganic phosphate on the mechanics and energetics of isometric contraction of fast skeletal muscle. Proceedings Biological sciences / The Royal Society 277: 19–27.

62 Potma, EJ, Stienen GJ. 1996. Increase in ATP consumption during shortening in skinned fibres from rabbit psoas muscle: effects of inorganic phosphate. J Physiol 496 (Pt 1): 1–12.

63 Potma EJ, van Graas IA, Stienen GJ. 1995. Influence of inorganic phosphate and pH on ATP utilization in fast and slow skeletal muscle fibers. Biophys J 69: 2580–9.

64 Amrute-Nayak, M, Antognozzi, M, Scholz, T, Kojima, H, et al. 2008. Inorganic phosphate binds to the empty nucleotide binding pocket of conventional myosin II. J Biol Chem 283: 3773–81.

65 Debold, EP, Beck, SE, Warshaw DM. 2008. Effect of low pH on single skeletal muscle myosin mechanics and kinetics. American journal of physiology Cell physiology 295: C173–9.

66 Tesi, C, Piroddi, N, Colomo, F, Poggesi C. 2002. Relaxation kinetics following sudden Ca(2+) reduction in single myofibrils from skeletal muscle. Biophys J 83: 2142–51.

67 Stehle, R, Kruger, M, Pfitzer G. 2002. Force kinetics and individual sarcomere dynamics in cardiac myofibrils after rapid ca(2+) changes. Biophys J 83: 2152–61.

68 Vilfan, A, Duke T. 2003. Instabilities in the transient response of muscle. Biophys J 85: 818–27.

69 Homsher, E, Wang, F, Sellers JR. 1992. Factors affecting movement of F-actin filaments propelled by skeletal muscle heavy meromyosin. Am J Physiol 262: C714–23.

70 Marston S. 2003. Random walks with thin filaments: application of in vitro motility assay to the study of actomyosin regulation. J Muscle Res Cell Motil 24: 149–56.

71 Rahman, MA, Salhotra, A, Mansson A. 2018. Comparative analysis of widely used methods to remove nonfunctional myosin heads for the in vitro motility assay. J Muscle Res Cell Motil 39: 175–87.

72 Nelson, CR, Debold, EP, Fitts RH. 2014. Phosphate and acidosis act synergistically to depress peak power in rat muscle fibers. American journal of physiology Cell physiology 307: C939–50.

73 Karatzaferi, C, Franks-Skiba, K, Cooke R. 2008. Inhibition of shortening velocity of skinned skeletal muscle fibers in conditions that mimic fatigue. American journal of physiology, Regulatory, integrative and comparative physiology 294: R948–55.

74 Forgacs, E, Sakamoto, T, Cartwright, S, Belknap, B, et al. 2009. Switch 1 mutation S217A converts myosin V into a low duty ratio motor. J Biol Chem 284: 2138–49.

75 Furch, M, Fujita-Becker, S, Geeves, MA, Holmes, KC, et al. 1999. Role of the salt-bridge between switch-1 and switch-2 of Dictyostelium myosin. J Mol Biol 290: 797–809.

76. Velayuthan LP, Moretto L, Tågerud S, Usaj M, et al. 2022. Virus-free, transfection, expression, and purification of human β-cardiac myosin using mouse myoblast cell line.; 49th European Muscle Conference (EMC)., Prague, Czech Republic.

77 Robert-Paganin, J, Auguin, D, Houdusse A. 2018. Hypertrophic cardiomyopathy disease results from disparate impairments of cardiac myosin function and auto-inhibition. Nature communications 9: 4019.

78 Parker, F, Peckham M. 2020. Disease mutations in striated muscle myosins. Biophysical reviews 12: 887–94.

79 Zananiri, R, Mangapuram Venkata, S, Gaydar, V, Yahalom, D, et al. 2022. Auxiliary ATP binding sites support DNA unwinding by RecBCD. Nature communications 13: 1806.

80 Amrute-Nayak, M, Lambeck, KA, Radocaj, A, Huhnt, HE, et al. 2014. ATP turnover by individual myosin molecules hints at two conformers of the myosin active site. Proc Natl Acad Sci U S A 111: 2536–41.

81 Heath, GR, Scheuring S. 2018. High-speed AFM height spectroscopy reveals micros-dynamics of unlabeled biomolecules. Nature communications 9: 4983.

82 Kovacs, M, Toth, J, Hetenyi, C, Malnasi-Csizmadia, A, et al. 2004. Mechanism of blebbistatin inhibition of myosin II. J Biol Chem 279: 35557–63.

83 Rauscher, AA, Gyimesi, M, Kovacs, M, Malnasi-Csizmadia A. 2018. Targeting Myosin by Blebbistatin Derivatives: Optimization and Pharmacological Potential. Trends Biochem Sci 43: 700–13.

84 Planelles-Herrero, VJ, Hartman, JJ, Robert-Paganin, J, Malik, FI, et al. 2017. Mechanistic and structural basis for activation of cardiac myosin force production by omecamtiv mecarbil. Nature communications 8: 190.

85 Kawas, RF, Anderson, RL, Ingle, SRB, Song, Y, et al. 2017. A small-molecule modulator of cardiac myosin acts on multiple stages of the myosin chemomechanical cycle. J Biol Chem 292: 16571–7.

86 Wigton, EJ, Thompson, SB, Long, RA, Jacobelli J. 2016. Myosin-IIA regulates leukemia engraftment and brain infiltration in a mouse model of acute lymphoblastic leukemia. J Leukoc Biol 100: 143–53.

87 Malik, FI, Hartman, JJ, Elias, KA, Morgan, BP, et al. 2011. Cardiac myosin activation: a potential therapeutic approach for systolic heart failure. Science 331: 1439–43.

88 Saberi, S, Cardim, N, Yamani, MH, Schulz-Menger, J, et al. 2020. Mavacamten Favorably Impacts Cardiac Structure in Obstructive Hypertrophic Cardiomyopathy: EXPLORER-HCM CMR Substudy Analysis. Circulation.

89 Desai, MY, Owens, AT, Geske, JB, Wolski, K, et al. 2022. Dose-Blinded Myosin Inhibition in Patients with Obstructive HCM Referred for Septal Reduction Therapy: Outcomes Through 32-Weeks. Circulation.

90 Gyimesi, M, Horvath, AI, Turos, D, Suthar, SK, et al. 2020. Single Residue Variation in Skeletal Muscle Myosin Enables Direct and Selective Drug Targeting for Spasticity and Muscle Stiffness. Cell 183: 335–46 e13.

